# *H. pylori* infection inhibits autophagy to aggravate DNA damage by p62-mediated Rad51 ubiquitination

**DOI:** 10.1101/771519

**Authors:** Chuan Xie, Nianshuang Li, Huan Wang, Cong He, Yi Hu, Chao Peng, Yaobin Ouyang, Dejie Wang, Yong Xie, Jiang Chen, Xu Shu, Yin Zhu, Nonghua Lu

## Abstract

*Helicobacter pylori* (*H. pylori*) infection is the strongest known risk factor for gastric carcinogenesis. DNA damage response (DDR) and autophagy play key roles in tumorigenic transformation. However, it remains unclear how *H. pylori* infection modulate DNA damage and autophagy. Here we report that *H. pylori* infection promotes DNA damage via suppression of Rad51 expression through inhibition of autophagy and accumulation of p62 in gastric carcinogenesis. We find that *H. pylori* infection caused alteration of DDR pathway and autophagy in gastric cells and Mongolian gerbils in a CagA-dependent manner. Moreover, loss of autophagy led to promotion of DNA damage in *H. pylori*-infected cells. Furthermore, knockdown of autophagic substrate p62 upregulated Rad51 expression, and p62 promoted ubiquitination of Rad51 via the direct interaction of the UBA domain with Rad51. Finally, *H. pylori* infection was associated with elevated levels of p62 in gastric intestinal metaplasia and decreased levels of Rad51 in dysplasia compared to their *H. pylori-* counterparts. Our findings provide a novel mechanism into the linkage of *H. pylori* infection, autophagy, DNA damage and gastric tumorigenesis.

## Introduction

*Helicobacter pylori (H. pylori)*, a human pathogen, colonizes approximately half of the world’s population, and its colonization can last for life without antibiotic intervention(Hooi, Lai et al., 2017, Zamani, Ebrahimtabar et al., 2018). *H. pylori* infection typically causes chronic inflammation and cellular injury, which leads to gastric carcinogenesis through the histopathological Correa cascade from chronic gastritis (CG), intestinal metaplasia (IM) and dysplasia (Dys) culminating in gastric cancer(Correa & Houghton, 2007, Kodaman, Pazos et al., 2014). Cytotoxin-associated gene A (CagA), encoded by the cag pathogenicity island, is a well-characterized virulence factor of *H. pylori*. CagA is delivered into gastric epithelial cells through the type IV secretion system, and dysregulates a series of intracellular signalling pathways, resulting in severe tissue damage and inflammation (Hatakeyama, 2014). It has been reported that CagA was associated with a higher risk of gastric cancer in experimental animals and human samples(Amieva & Peek, 2016).

The maintenance of human genome integrity depends on DNA damage response. Although single-strand breaks (SSB) is the most common types of DNA damage, double-strand breaks (DSB) is lethal to cells and need to be rapidly repaired for ensuring cell survival. In response to DSB, DNA damage sensor Ataxia-telangiectasia mutated serine/threonine kinase (ATM) is rapidly translocated to the sites of DNA damage where it is activated(Blackford & Jackson, 2017, Scully, Panday et al., 2019). The activation of ATM phosphorylates histone H2AX (γH2AX) and induces damage-pathway downstream effectors, such as Chk2, p53 and BRCA1, leading to DNA repair, cell cycle arrest and apoptosis(Huen & Chen, 2008). The highly conserved protein Rad51 plays a central role in homologous recombination of DNA during DSB repair. It has been shown that accumulation of DNA damage by elevated mutation rate, a potential cause of cancer, is associated with altered Rad51 expression(Gachechiladze, Skarda et al., 2017, Lose, Lovelock et al., 2006). Accumulating evidence has indicated that *H. pylori* infection increases production of reactive oxygen species (ROS) and DNA damage levels, which can contribute genome instability and gastric carcinogenesis(Xie, Yi et al., 2018). Also, our previous studies indicated a progressively elevated γH2AX levels in gastric pathological cascade from CG, IM to Dys, but slightly declined in GC, suggesting that DSBs appear to be an early event in *H. pylori*-mediated genetic instability and gastric tumorigenesis(Xie, Xu et al., 2014). Although DNA damage can be efficiently repaired, it has been reported that long-term bacterial infection leads to accumulation of unrepaired breaks due to saturation of repair capabilities (Toller, Neelsen et al., 2011). However, the molecular mechanism underlying *H. pylori*-induced DSBs and the DNA damage repair has not been fully elucidated.

Recently, DNA damage has been linked to autophagy, an intracellular degradation process by which cytoplasmic materials are delivered to the lysosome for digestion(Eliopoulos, Havaki et al., 2016). In response to various cellular stress, activation of autophagy is crucial for maintaining cellular energy levels through synthesis, degradation and recycle of constituents. Induction of autophagy serves as a cell defence mechanism against intracellular pathogenic microorganisms(Sudhakar, Jacomin et al., 2019). Several microbes such as Listeria monocytogenes have been shown to stimulate formation of autophagosomes, which in turn initiates bacterial degradation(Huang & Brumell, 2014). Emerging evidence over the past decade has indicated that *H. pylori* infection was linked to autophagy(Horvat, Noto et al., 2018). The virulence factor cytotoxin (VacA) is sufficient to induce autophagy and cell death in *H. pylori*-infected gastric epithelial cells(Zhu, Xue et al., 2017). Furthermore, another important bacterial oncogenic protein CagA has been found to be degraded by autophagy, whereas CAPZA1 overexpression promoted CagA to escape from autophagic degradation(Tsugawa, Mori et al., 2018). In addition, *H. pylori* CagA protein was reported to negatively regulate autophagy and promote inflammation(Li, Tang et al., 2017).

Activation of autophagy by various cellular stress has been implicated in DNA repair mechanisms, such as homologous recombination (HR) and non-homologous end-joining (NHEJ)(Hewitt & Korolchuk, 2017). Degradation of p62/SQSTM1 (p62), the best-known autophagic substrate, has been widely identified as an indicator of autophagy. p62 interacts with ubiquitinated proteins via its C-terminal ubiquitinated-associated domain (UBA) and results in their autophagic degradation(Peng, Yang et al., 2017). Wang et al. have found that p62 accumulation due to deficiency of autophagy can bind to and inhibit the activity of RNF168, an E3 ligase required for histone H2A ubiquitination. As a result, DSB repair pathway was significantly impaired(Wang, Zhu et al., 2017). In addition, overexpression of p62 has been shown to induce proteasomal degradation of homologous recombination marker Rad51(Hewitt, Carroll et al., 2016). Currently, the precise mechanisms by which autophagy mediates DDR and genome stability remains unknown in *H. pylori* infection-associated gastric tumorigenesis.

In this study, we show that prolonged exposure to *H. pylori* infection inhibited autophagosome formation, which contributed to excessive DNA damage and eventual gastric tumorigenesis. Mechanistically, accumulation of p62 due to autophagy inhibition caused DNA repair marker Rad51 ubiquitination and degradation. Our findings indicated a critical role for autophagy and DDR in *H. pylori* infection-induced gastric carcinogenesis.

## Results

### *H. pylori* infection promote ATM-dependent DNA damage response

We have previously shown that γH2AX, a reliable marker of DSB, was progressively increased in tissues in gastric lesions from CG, IM to Dys(Xie et al., 2014).Given an important role of *H. pylori* infection in gastric carcinogenesis, we investigated whether *H. pylori* infection induces DNA damage in gastric epithelial cells. Human normal gastric epithelial GES-1 cells were co-cultured with *H. pylori* strain 43504 at various time-points and at different MOIs. γH2AX was significantly increased in a time- and MOI-dependent manner following infection (Figure 1A and 1B). In addition, a percentage of tail DNA in the comet assay was used to represent the abundance of fragmented DNA due to DNA damage. In GES-1 cells, *H. pylori* infection led to significant increases of DNA DSB levels over the infection duration (Figure 1C). To further evaluate these in vitro findings, Mongolian gerbil, a widely used model for investigating *H. pylori*-induced gastric tumorigenesis(Peng, Li et al., 2019), were challenged with Brucella broth (negative control) or with carcinogenic *H. pylori* 43503 strain. Consistent with our in vitro data, *H. pylori* infection significantly increased gastric γH2AX expression levels compared to uninfected control and elevated γH2AX expression at 12 months post-infection (MPI) compared to 6 MPI using immunohistochemistry (Figure 1D). It has been documented that ATM is recruited and activated at sites of DSB where it phosphorylates serine/threonine kinase Chk2 at priming site Thr68, leading to cell cycle arrest through phosphorylation of p53(Roos, Thomas et al., 2016). We next examined whether *H. pylori*-induced DNA damage is associated with the activation of ATM. Following *H. pylori* infection on gastric GES-1 cells, levels of phosphorylated ATM were increased in a time-dependent manner. In addition, its downstream substrates Chk2 and p53 were also activated via phosphorylation (Figure 1E). Furthermore, the percentage of S phase was significantly increased in gastric epithelial cells at 12h or 24h of infection, suggesting that *H. pylori* could induce S-phase cell cycle arrest (Figure 1F).

**Figure 1.**
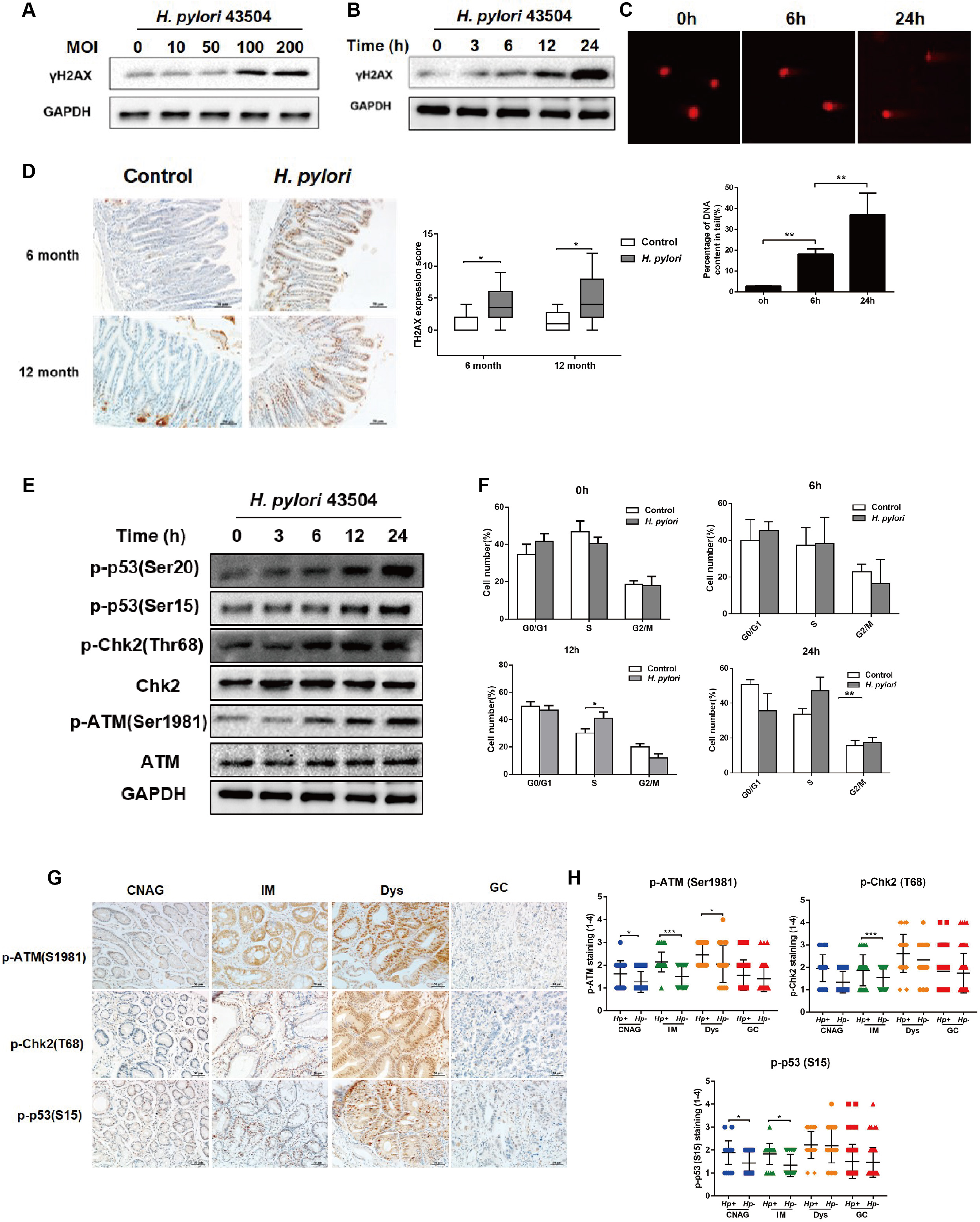
*H. pylori* infection induces DNA damage response signalling pathway. (A, B) Western blot analysis of γH2AX in GES-1 cells with *H. pylori* 43504 infection (A) at different MOI or (B) at different time points. (C) The comet assay for assessing DNA damage in GES-1 cells with or without *H. pylori*. (D) Immunochemistry for γH2AX in gastric tissues collected from Mongolian gerbils that were challenged with *H. pylori* 43504 strain or Brucella broth. (E) Western blot analysis of p-p53 (Ser20), p-p53 (Ser15), p-Chk2 (Thr168), Chk1, p-ATM (Ser1981), ATM in GES-1 cells infected with *H. pylori* 43504 at different time points. (F) Flow cytometry analysis for cell cycle distribution after *H. pylori* infection. (G) Representative immunohistochemical staining for p-ATM (S1981), p-Chk2 (T68) and p-p53 (S15) in a series of human gastric tissues with Correa cascade from CNAG, IM, Dys and GC. (H) Quantitative analysis of the immunohistochemical staining results in gastric tissues. \**P*<0.05, \*\**P*<0.01.

To characterize the role of the DNA damage response pathways in *H. pylori*-associated gastric carcinogenesis, we examined and compared the protein levels of phosphorylated ATM, Chk2 and p53 in human gastric tissues with CG, IM to Dys and between *H. pylori*-positive and -negative tissues. Levels of p-ATM (S1981), p-Chk2 (T68) and p-p53 (S15) were gradually increased during neoplastic progression from CG, IM to Dys. *H. pylori* infection induced the significant increases of p-ATM at CG/IM/Dys, p-Chk2 at IM and p-p53 at CG/M compared to the *H. pylori*-negative subjects (Figure 1H). Interestingly, the activity of these protein targets was decreased in GC stage (Figure 1G and 1H). In addition, there was no significant differences on total ATM, Chk2 levels in human gastric tumorigenesis cascade (Figure S1A and S1B). Taken together, our findings clearly indicate that *H. pylori* infection promoted the DNA damage response pathway in the early stages of gastric tumorigenesis.

### Persistent *H. pylori* infection promoted DNA damage via suppression of Rad51

Accurate DNA damage repair is required for maintaining genome stability. Our previous studies have shown that H. pylori infection was correlated with the expression of NHEJ marker Ku70/80 in human GC tissues(Li, Xie et al., 2013). To explore how *H. pylori* infection modulates DSB HR repair, GES-1 cells were co-cultured with *H. pylori* 43504 strain at different time-points. *H. pylori* infection resulted in the reduction of Rad51 protein, a crucial protein in HR DNA repair process, but did not significantly affect another DNA repair marker BRCA2, which can facilitate the loading of the recombination protein Rad51 at DNA breaks (Figure 2A)(Jensen, Carreira et al., 2010). In addition, *H. pylori* infection led to statistically significant increase in the ratio of γH2AX and Rad51. γH2AX and Rad51 focus formation were detected by immunofluorescent microscopy (Figure 2B). These findings suggested that *H. pylori*-induced elevation of DSBs may be due to downregulation of Rad51. To further elucidate the role of Rad51 in DSB repair, gastric GES-1 cells were treated with Rad51-expressing plasmid or Rad51 siRNA and then co-cultured with *H. pylori* 43504 strain. *H. pylori*-induced elevation of γH2AX and downstream phosphorylated p53 levels were suppressed in cells transfected with the Rad51 plasmid (Figure 2C). Conversely, siRNAs-mediated Rad51 knockdown led to an increase of levels of γH2AX and phosphorylated p53 (Figure 2D). Moreover, overexpression of Rad51 significantly suppressed *H. pylori*-induced DNA damage, whereas inhibition of Rad51 upregulated DNA damage as demonstrated in the comet assay, further indicating the importance of Rad51 in *H. pylori*-induced DNA damage response (Figure 2E). To evaluate if these in vitro mechanisms are operable in vivo, Rad51 levels in the gastric mucosa of Mongolian gerbils at 6 and 12 MPI were examined and quantified using immunohistochemistry analysis. At 6 MPI, elevated Rad51 levels were found between infected and control groups, but there was no statistic difference. By 12 MPI, Rad51 expression was significantly decreased in the *H. pylori*-infected gerbils versus the control animals (Figure 2F and 2G). Collectively, these data indicate that persistent *H. pylori* infection suppressed expression of homologous recombination marker Rad51, which resulted in the accumulation of DNA damage.

**Figure 2.**
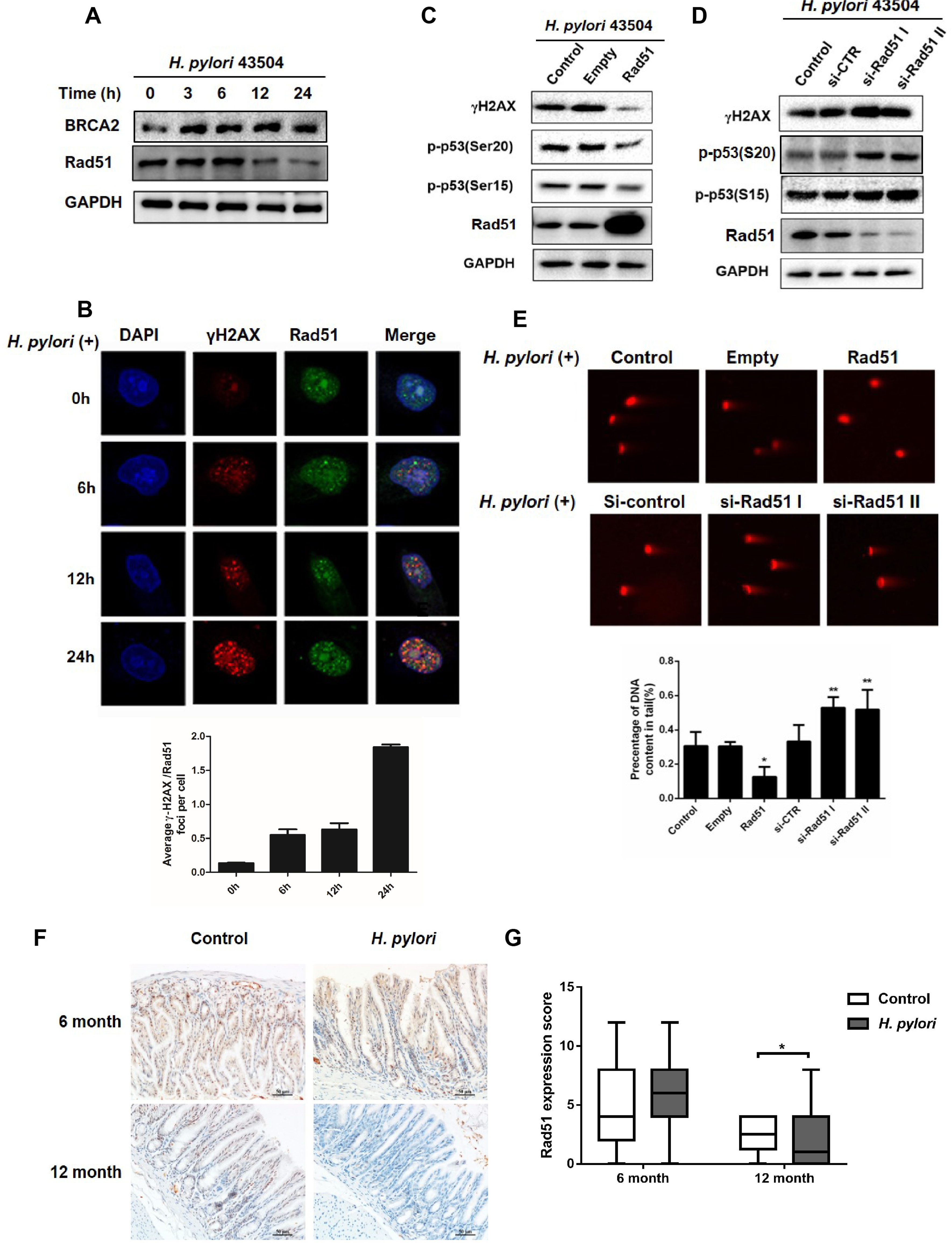
*H. pylori* infection cause persistent DNA damage through inhibition of Rad51. (A) Western blot analysis of Rad51 in GES-1 cells with *H. pylori* 43504 infection. (B) Immunofluorescence staining of γH2AX (Red) and Rad51 (Green) in gastric cells with *H. pylori* infection at different time. DAPI counterstaining was shown in blue. (C, D) Western blot analysis of γH2AX, p-p53 (Ser20) and p-p53 (Ser15) in GES-1 cells, (C) which were treated with *H. pylori* in combination with plasmid expressing Rad51, (D) which were infected with *H. pylori* and transfected with Rad51 siRNAs. (E) After transfected with Rad51 expression plasmid or Rad51 siRNAs, the comet assay was performed to detect DNA damage and the length of tail DNA was evaluated in gastric cells with *H. pylori* infection. (F) Immunohistochemistry staining for Rad51 in gastric tissues of Mongolian gerbils infected with *H. pylori*. Scale bar: 50μm. \**P*<0.05, \*\**P*<0.01.

### Autophagy are dynamically changed in response to *H. pylori* infection

Autophagy, a cellular self-degradation process, plays a pivotal role in host-pathogen interaction and gastric carcinogenesis. LC3B is a well-established autophagy marker. Conversion of soluble LC3B-I to lipid bound LC3B-II has been considered as a hallmark of autophagosome formation (Pugsley, 2017). To investigate how *H. pylori* infection affects autophagy, gastric cells were co-cultured with *H. pylori* at different time-points. Western blot showed that levels of LC3B-I and II were significantly increased up to 6 hours post-infection (HPI) and then were gradually decreased (Figure 3A). Consistently, transmission electron microscopy (TEM) revealed that there was an increase of autophagosomes or autolysosomes-like structures at 6 HPI, which were significantly reduced by 24 HPI (Figure 3B). P62, a well-known substrate for autophagic degradation, can be used as an indicator of autophagic flux(Moscat, Karin et al., 2016). As shown in Figure 3C and 3D, p62 levels in the gastric mucosa of *H. pylori*-infected gerbils were significantly increased at 12 MPI.

**Figure 3.**
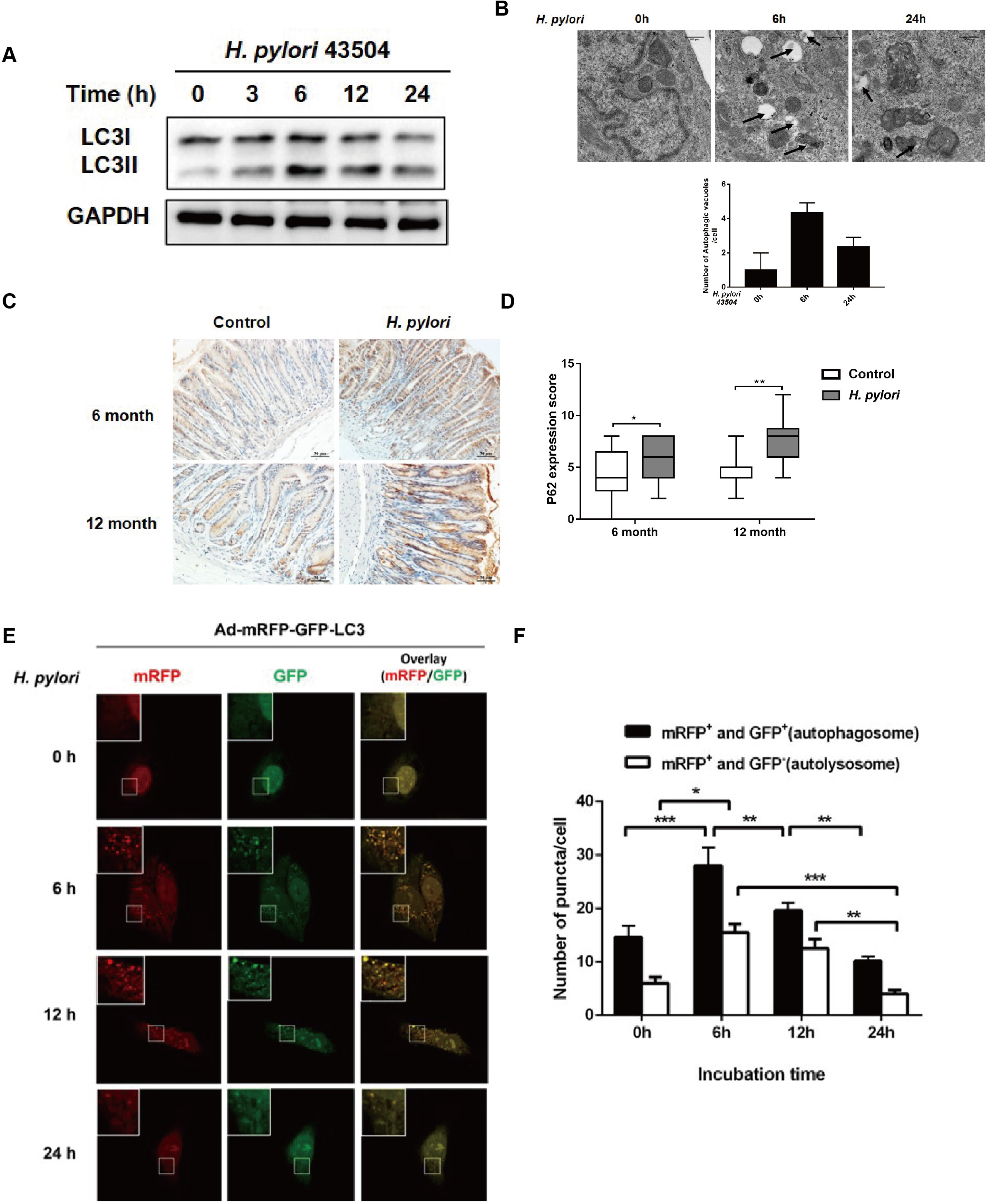
Autophagy are dynamically changed in response to *H. pylori* infection. (A) Western blot analysis of LC3 in GES-1 cells co-cultured with *H. pylori* 43504. (B) Transmission electron microscopy showing autophagosomes in gastric cells with or without *H. pylori* infection. Scale bar: 500μm. (C) Representative p62 immunohistochemical staining in gastric tissues. Scale bar: 50μm. (D) Quantification of staining scores for p62 in gastric epithelium. (E) After transfection with mRFP-GFP-LC3 adenovirus, cells were incubated with *H. pylori* and Confocal microscopy was used to monitor the autophagic flux. (F) The numbers of autophagosome (mRFP^+^GFP^+^) and autolysosome (mRFP^+^GFP^-^) were quantified. \**P*<0.05, \*\**P*<0.01, \*\*\**P*<0.001.

To directly visualize and quantitatively characterize autophagy *in situ*, adenovirus expressing LC3B fused to fluorescent proteins mRFP and GFP (Ad-mRFP-GFP-LC3) was transfected into GES-1 cells in which, autophagosome formation with the mRFP-positive/GFP-positive (yellow) puncta can be distinguished from autolysosome with the mRFP-positive/GFP-negative puncta (red). The number of autophagosomes and autolysosomes in the cells were increased at 6 HPI. However, by 24 HPI, both the abundance of autophagosomes and autolysosomes were rapidly decreased (Figure 3E and 3F). Taken together, these findings indicate that *H. pylori* infection induces autophagy at 6 HPI, which was followed by a rapid decrease of autophagy by 24 HPI.

### Inhibition of autophagy increased DNA damage in response to *H. pylori* infection

Recent studies have documented that cell autophagy contributes to the maintenance of the genomic stability and loss of autophagy impairs DNA repair(Eliopoulos et al., 2016). We next investigated whether autophagy plays a role in DNA damage response to infection in gastric GES-1 cells. GES-1 cells were treated with the autophagy inhibitor 3-MA, or BafA1, the autophagy inducer rapamycin, or starvation, and then co-cultured with *H. pylori* strain 43504. Inhibition of autophagy by the pharmacological inhibitors significantly aggravated the *H. pylori*-induced DNA damage signalling pathway, which were reflected by the upregulation of DNA damage marker γH2AX, increased the phosphorylation of p53 and the downregulation of DNA repair protein Rad51, and opposite is true with the treatment of rapamycin or starvation that activates autophagy (Figure 4A). These results were further supported using immunofluorescence. (Figure 4B). In addition, the tail lengths of *H. pylori*-induced-DNA fragmentation were significantly increased in response to the treatment of 3-MA or BafA1, but decreased after treatment with rapamycin or starvation (Figure 4C and 4D). Furthermore, *H. pylori* infection concurrently with the treatment of the autophagy activator Rapamycin or nutrient starvation accelerated cell cycle progression from S to G2/M phase comparted to *H. pylori* infection alone (Figure 4E). Taken together, these data suggested that persistent *H. pylori* infection led to the accumulation of DNA damage and cell cycle arrest via the suppression of autophagy.

**Figure 4.**
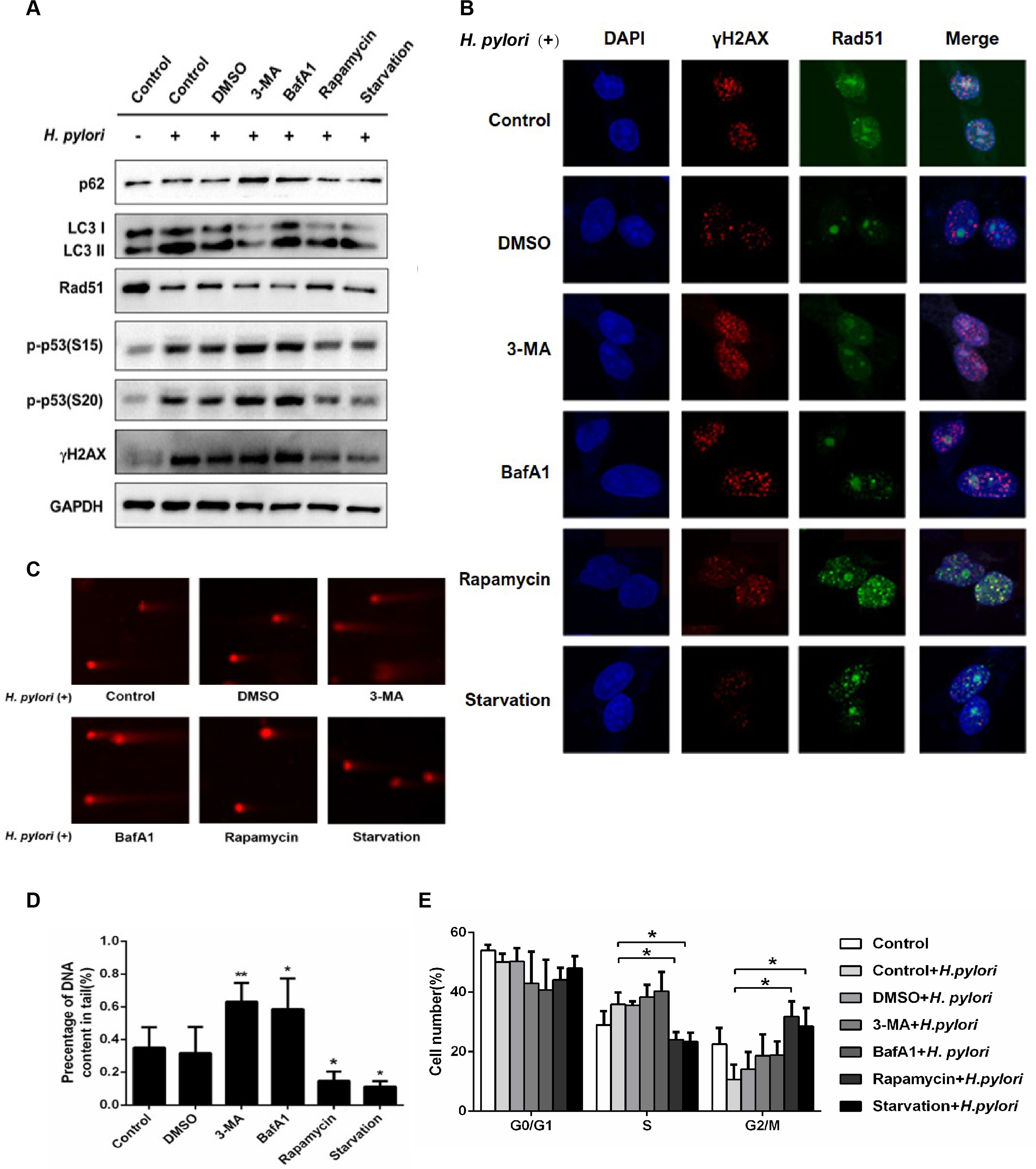
Inhibition of autophagy increased DNA damage in response to *H. pylori* infection. (A) Western blot analysis of p62, LC3, Rad51, p-p53 (S15), p-p53 (S20), γH2AX protein following *H. pylori* infections and treatment with autophagy inhibitors (3-MA, BafA1) or autophagy activators (Rapamycin, Starvation). (B) Immunofluorescence assay showing γH2AX and Rad51 expressions in GES-1 cells treated with autophagy inhibitors or autophagy activators and then infected with *H. pylori*. (C) After treatment with autophagy related drugs and *H. pylori*, DNA damage was determined by the Comet assay in GES-1 cells. (D) Comet tail lengths showing DNA damage. (E) Flow cytometry analysis showing cell cycle progression in cells treated with autophagy inhibitor or activators and then infected with *H. pylori*. \**P*<0.5, \*\**P*<0.01.

### Rad51 is integral for autophagy-mediated DNA damage repair response to *H. pylori* infection

Our aforementioned results indicated that *H. pylori* infection suppressed autophagy and DNA repair protein Rad51. Additionally, inhibition of autophagy led to decreased expression of Rad51 in the setting of *H. pylori* infection. We hypothesize that Rad51 could serve as a crucial adaptor in autophagy-regulated DNA damage in response to *H. pylori* infection. To test this hypothesis, GES-1 cells were transfected with Rad51-expressing plasmid, in combination with *H. pylori* 43504 infection and treatment of autophagy inhibitors 3-MA or BafA1. Overexpression of Rad51 suppressed elevation of *H. pylori*-induced DNA damage signals such as γH2AX and its downstream factors due to impairment of autophagy (Figure 5A and 5B). Conversely, knocking-down endogenous Rad51 by siRNA significantly increased *H. pylori*-induced DNA damage signal suppressed by the activation (rapamycin or starvation) of autophagy (Figure 5C and 5D). Consistent with the Western blot data, transient overexpression of Rad51 decreased the *H. pylori*-induced DNA fragmentation, despite of inhibition of autophagy, whereas, the knockdown of Rad51 elevated autophagy-inhibited DNA damage (Figure 5E and 5F). Collectively, these data suggest that autophagy modulates *H. pylori*-induced DNA damage via Rad51.

**Figure 5.**
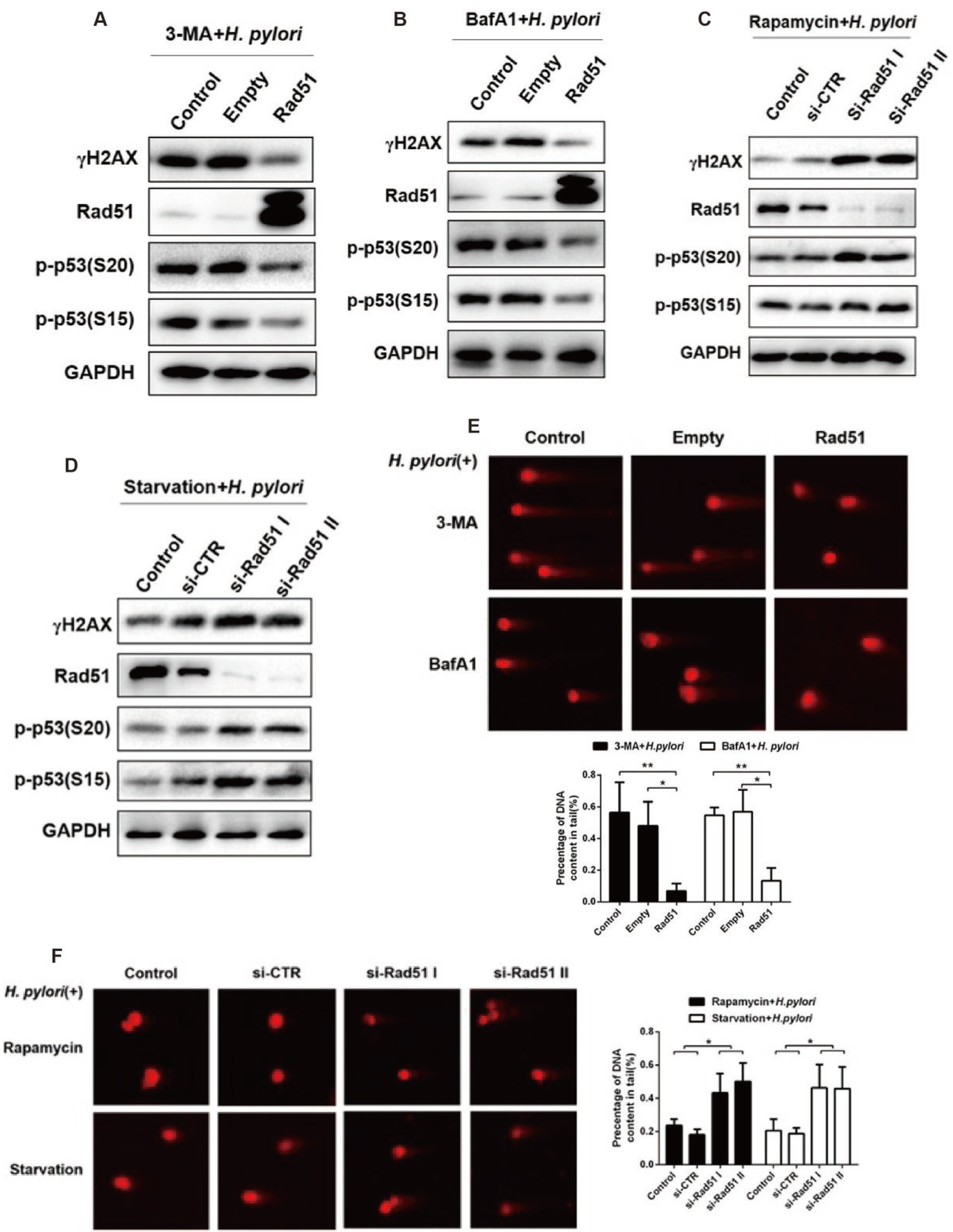
Rad51 is a crucial adaptor in autophagy regulated DNA damage in response to *H. pylori* infection. (A, B) Western blot analysis of γH2AX, Rad51, p-p53 (S20) and p-p53 (S15) protein levels in GES-1 cells following *H. pylori* infection, and transfection with Rad51 expression plasmids in the presence of autophagy inhibitor treatment (A) 3-MA or (B) BafA1. (C, D) Western blot showing expression of γH2AX, Rad51, p-p53 (S20) and p-p53 (S15) in GES-1 cells transfected with Rad51 siRNAs, infected with *H. pylori* and treated with autophagy inducer (C) rapamycin or (D) starvation. (E) Comet assay showing DNA damage in GES-1 cells transfected with Rad51 expression plasmid, and then infected with *H. pylori* in the presence of autophagy inhibitors 3-MA or BafA1. (F) Comet assay showing DNA damage in GES-1 cells transfected with Rad51 siRNAs, and then infected with *H. pylori* in the presence of autophagy inducers rapamycin or starvation. \**P*<0.5, \*\**P*<0.01.

### *H. pylori*-induced DNA damage response and suppression of autophagy are cytopathic events common to different CagA^+^*H. pylori* strains, cell types and animal models

To examine whether *H. pylori*-induced DNA damage response and alteration of autophagy are strain-specific. CagA^+^*H. pylori* strain 7.13 was co-cultured with GES-1 cells with different MOI for 24h. Consistent with the results with ATCC43504 (Figure 1A and 2A), the strain 7.13 infection in GES-1 cells increased levels of the DSBs marker γH2AX in concert with the decrease of the DNA repair protein Rad51 in an MOI-dependent manner. Also, accumulation of autophagy substrate p62 ascended in line with an increase of MOIs following 7.13 infection (Figure 6A). TEM assay also showed that 7.13 infection increased the number of autophagosomes at 6 HPI, and then led to the inhibition of autophagy at 24 HPI. These findings with 7.13 strain was consistent with the results with ATCC43504 strain (Figure 6B). To exclude a possibility that *H. pylori*-induced cytopathic effects were cell type-dependent, gastric adenocarcinoma AGS cells were co-cultured with 7.13 strain. The responses of the DNA damage marker γH2AX, DNA repair protein Rad51 and the autophagy substrate p62 to 7.13 infection in AGS cells were similar to those in GES-1 cells (Figure 6C). CagA protein is one of the most important virulence factors in *H. pylori*-induced gastric carcinogenesis. We therefore investigated the role of CagA in DNA damage and autophagy regulation. AGS and GES-1 cells were co-cultured with *H. pylori* strain 7.13 or its isogenic CagA-mutant. Unlike wild-type *H. pylori* 7.13, the loss of CagA failed to promote DNA damage and accumulation of autophagic receptor p62, and to suppress expression of Rad51 (Figure 6D).

**Figure 6.**
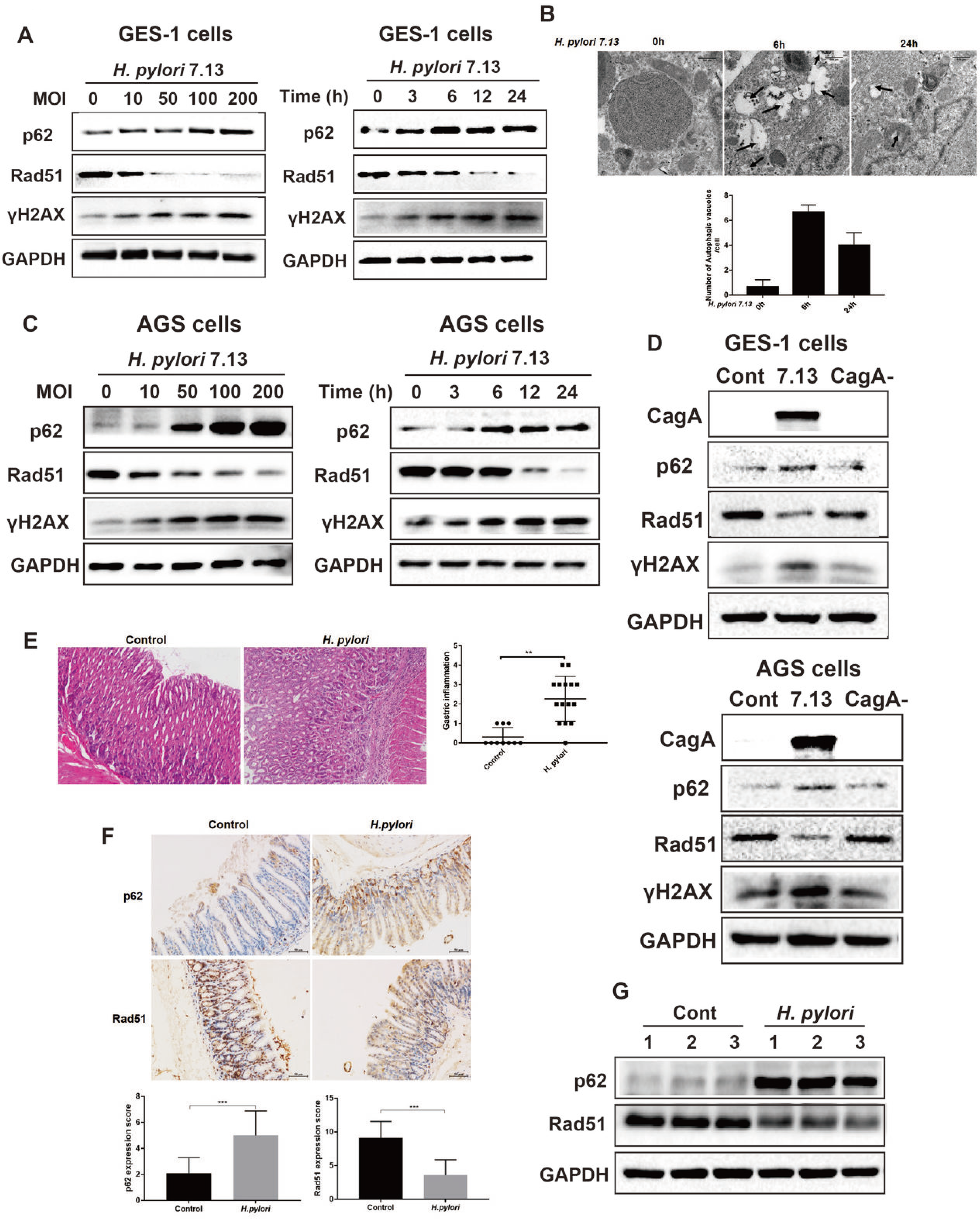
*H. pylori*-Susceptible DNA damage response and autophagy are shared between different bacterial strains, cell types and animal models. (A) Western blot of p62, Rad51 and γH2AX in GES-1 cells infected with *H. pylori* 7.13 at different MOI or at different time points. (A) Gastric cancer cells were infected with *H. pylori* 7.13 at different MOI or at different time points, and then immunoblot assay was performed to assess the expression of p62, Rad51, and γH2AX. (B) Transmission electron microscopy for autophagic vacuoles in gastric cells infected with *H. pylori* 7.13. Scale bar: 500μm. (C, D) Western blot analysis showing the expression of CagA, p62, Rad51 and γH2AX in gastric cells (C) GES-1, or (D) AGS infected with *H. pylori* strain 7.13 or its CagA-knockout mutant. (E) H&E staining for gastric pathology in gastric tissues of *H. pylori*-infected C57BL/6 mice. (F) Immunohistochemistry staining for Rad51 and p62 levels in C57BL/6 mice infected with *H. pylori*. Scale bar: 50μm. (G) Western blot analysis of Rad51 and p62 protein levels in gastric tissues of *H. pylori*-infected mouse model. \*\**P*<0.01, \*\*\**P*<0.001.

To explore whether *H. pylori* infection-regulated DNA damage and autophagy responses are operable in different animal models, C57BL/6 male mice were infected with *H. pylori* 43504 strain for 3 months. *H. pylori* colonization of mouse gastric mucosa was visualized using immunohistochemistry and FISH (Figure S2A and S2B). Real-time PCR analysis showed that infection with *H. pylori* led to transcriptional upregulation of inflammatory factors including *IL-1β*, *IL-6*, *IL-10* and *IL-17*, suggesting inflammation response induced by *H. pylori* (Figure S2C). H&E staining showed that *H. pylori* infection resulted in significantly gastric inflammation compared to control group at 3MPI (Figure 6E). In addition, levels of Rad51were significantly decreased in concert with increased p62 levels in *H. pylori*-infected gastric mucosa, compared to uninfected controls (Figure 6F). This result was supported by Western blot analysis showing the reduction of Rad51 level and the elevation of p62 level in *H. pylori*-infected mouse gastric mucosa (Figure 6G). These results demonstrate that accumulation of DNA damage and loss of autophagy due to *H. pylori* infection were operable by other CagA^+^*H. pylori* strains, in different cell lines or different animal models.

### Accumulation of p62 promoted *H. pylori*-induced DNA damage via suppression of the DNA repair protein Rad51

Given that *H. pylori*-induced suppression of autophagy was linked to persistent DNA damage and genome instability via inhibition of DNA repair protein Rad51, we next investigated the interaction between autophagy and Rad51. P62 is the best-known autophagic substrate. We hypothesize that p62 is a crucial factor in modulating expression of Rad51 since levels of p62 was inversely related with levels of Rad51 levels in *H. pylori*-infected gastric cells and gastric mucosa of animal models. Overexpression of p62 in AGS or GES-1 cells significantly reduced the expression of Rad51 (Figure 7A). Whereas knockdown of p62 led to upregulation of Rad51 protein (Figure 7B). To further assess how H. pylori-induced accumulation of p62 regulates expression of Rad51. Normal gastric epithelial GES-1 cells were transfected with p62 shRNA plasmid and cocultured with or without *H. pylori* 7.13 strain. The knockdown of p62 with the p62 shRNA plasmid increased the expression of Rad51 in both *H. pylori* 7.13-infected GES-1 and AGS cells (Figure 7C and 7D). Furthermore, we assessed whether p62 also suppressed Rad51 expression in *H. pylori* 43504-infected C57BL/6 mice in combination with treatment of everolimus, a well-known autophagy inducer of inhibiting mTOR. As shown in Figure 7E, everolimus treatment significantly attenuated H. pylori-induced inflammatory response at 3 MPI (Figure 7E). Immunohistochemistry staining revealed that the gastric mucosa of C57BL/6 mice treated with both *H. pylori* infection and everolimus contained lower p62 levels and higher Rad51 levels, compared to the mice infected with *H. pylori* alone (Figure 7F). The loss of p62 caused by everolimus was significantly correlated with the elevation of Rad51 levels following *H. pylori* infection (Figure 7G). These results indicate that accumulation of the autophagic substrate p62 in *H. pylori*-infected gastric epithelial cells promoted *H. pylori*-induced DNA damage via suppression of the DNA repair marker Rad51.

**Figure 7.**
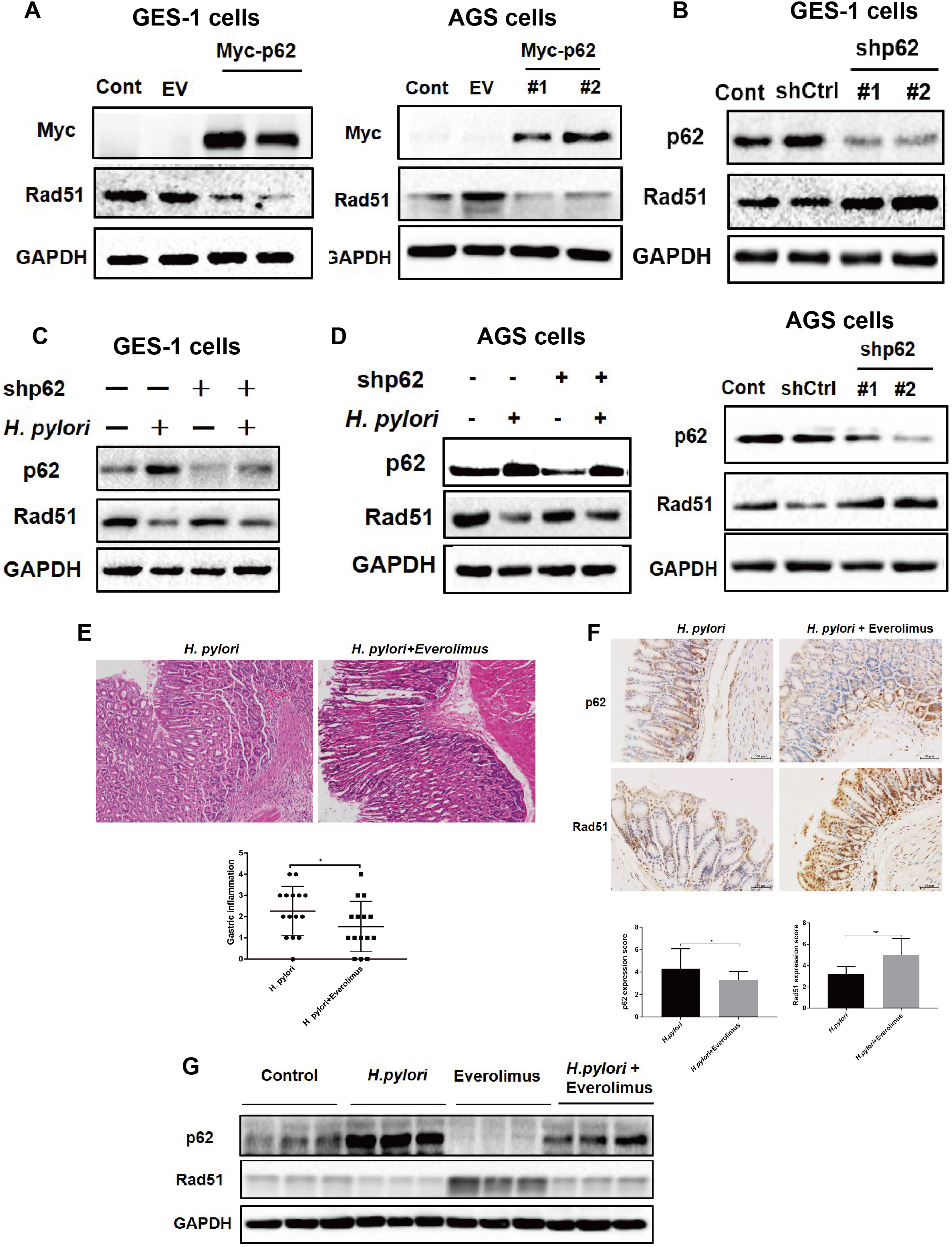
Accumulation of p62 suppressed DNA repair protein Rad51 in response to *H. pylori* infection. (A) Western blot analysis of p62 and Rad51 proteins in GES-1 and AGS cells transfected with plasmid expressing p62. (B) Western blot analysis of p62 and Rad51 proteins in GES-1 and AGS cells transfected with p62 shRNAs. (C, D) After transfected with p62 shRNAs, (C) GES-1 cells or (D) AGS cells showing p62 and Rad51 levels by immunoblot assay following *H. pylori* infection. (E) H&E staining showing the extent of gastric inflammatory response in C57BL/6 mice infected with *H. pylori* alone or combined with everolimus treatment. (F) Immunohistochemistry staining showing p62 and Rad51 levels in C57BL/6 mice treated with *H. pylori* and autophagy inducer everolimus by gastric gavage. Scale bar: 50μm. (G) Western blot analysis for p62 and Rad51 in gastric tissues of mouse models.

### The ubiquitination associated (UBA) domain of p62 was essential for *H. pylori* infection-mediated Rad51 ubiquitination

To test whether *H. pylori* infection induced proteasomal degradation of Rad51, AGS cells were pre-treated with a proteasome inhibitor MG132 prior to *H. pylori* 7.13 infection. Inhibition of proteasomal function by MG132 led to an increase of Rad51 levels (Figure 8A). In addition, AGS or GES-1 cells were co-transfected with plasmids expressing Rad51 and ubiquitin, then treated with proteasomal inhibitor MG132 for 6 hours prior to *H. pylori* 7.13 or its CagA^-^ mutant infection. CagA^+^ *H. pylori* 7.13 but not its strain its CagA^-^ mutant significantly promoted Rad51 ubiquitination (Figure 8B). These results indicate that CagA^+^*H. pylori* infection decreased Rad51 through the ubiquitination and degradation of Rad51. Next, we dissected the mechanism underlying *H. pylori*-induced accumulation of p62 and suppression of Rad51. Our data showed that *H. pylori* infection suppressed Rad51 protein expression, but not mRNA, suggesting that post-transcriptional process affected Rad51 protein levels (Figure S3A). Given that p62, an ubiquitin and LC3 binding protein, directly binds to an ubiquitinated protein via its UBA domain and sequesters them into inclusion for degradation by autophagy(Lee & Weihl, 2017), we examined the possible interaction between p62 and Rad51. Immunofluorescence staining showed that p62 were mostly localized in the cytoplasm and nucleus of uninfected cells, respectively. *H. pylori* infection could enhance p62 levels and decreased Rad51 expression. Furthermore, *H. pylori* infection resulted in the nuclear translocation of p62 where p62 was colocalized with Rad51, suggesting that *H. pylori* infection promoted p62 nuclear entry and interaction with Rad51 (Figure 8C). To demonstrate the physical interaction between p62 and Rad51, Plasmids Flag-Rad51 or Myc-p62 were transfected into stomach AGS or GES-1 cells followed by the immunoprecipitation assay. As shown in Figure 8D, p62 were coimmunoprecipitated with could interact with Rad51, indicating that p62 interacted with Rad51(Figure 8D).

**Figure 8.**
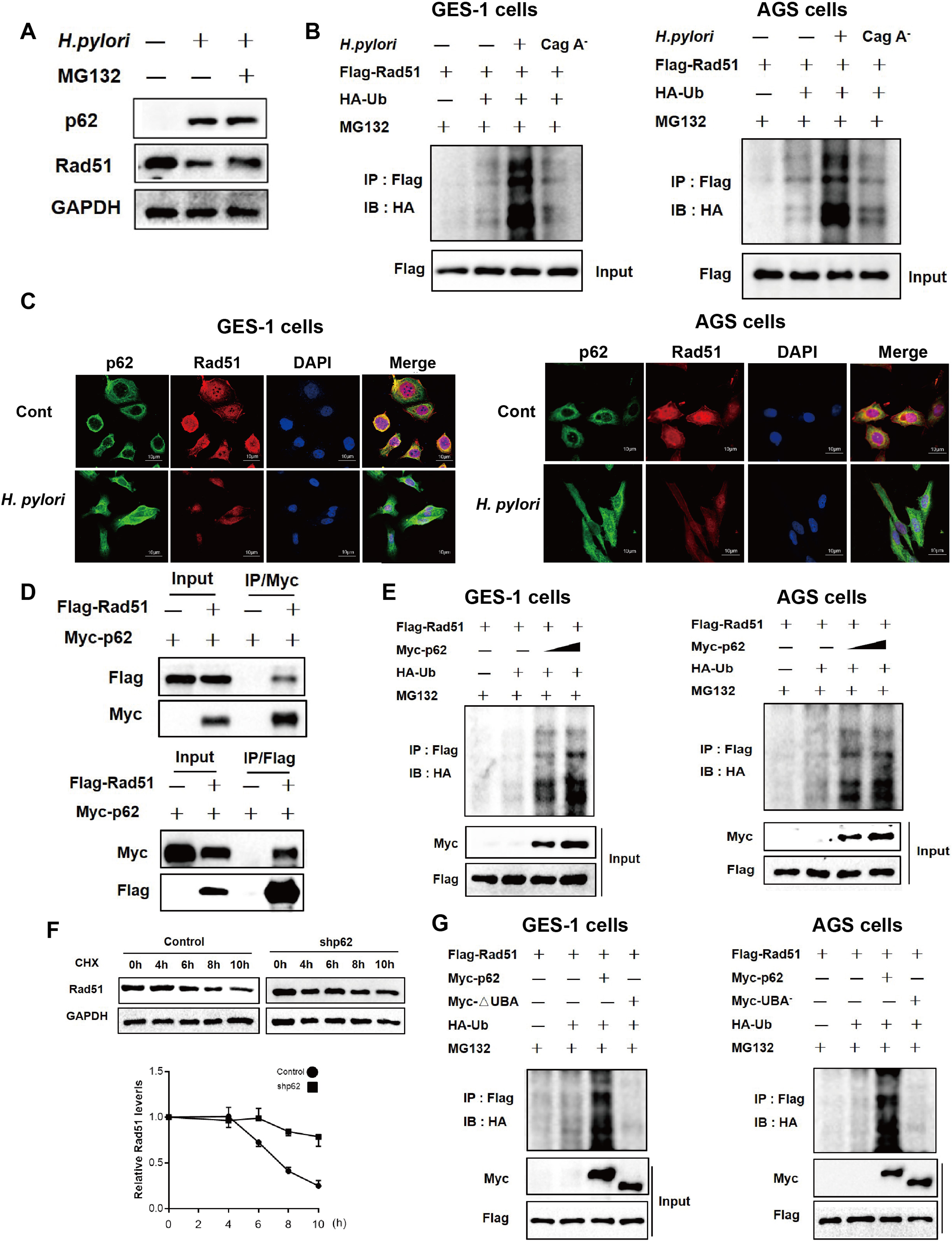
P62 induced *H. pylori* infection-mediated Rad51 ubiquitination. (A) Gastric epithelial cells were co-cultured with *H. pylori* strain and incubated with proteasome inhibitor MG132. P62 and Rad51 protein levels were measured by Western blot assay. (B) AGS or GES-1 cells were co-transfected with plasmids expressing Rad51 and ubiquitin, then infected with *H. pylori* strain 7.13 or its CagA-knockout mutant and incubated with proteasome inhibitor MG132. Lysates were detected for polyubiquitinated Rad51 by immunoprecipitation assay. (C) Immunofluorescence assay for the intracellular localization of Rad51 and p62 in gastric epithelial cells following *H. pylori* infection. (D) The interaction between Myc-p62 and Flag-Rad51 was examined using immunoprecipitation assay. (E) AGS or GES-1 cells were co-transfected with plasmids of Flag-Rad51, Myc-p62 and HA-ubiquitin, and then incubated with MG132. Immunoprecipitation assay was used to determine polyubiquitinated Rad51. (F) Stability of Rad51 protein was analysed in gastric epithelial cells transfected with p62 shRNA using the cycloheximide chase method. (G) AGS or GES-1 cells were co-transfected with plasmids of Flag-Rad51, Myc-p62, Myc-UBA^-^ and HA-ubiquitin, and then incubated with MG132. Immunoprecipitation assay was used to determine polyubiquitinated Rad51

To determine the role of p62 in modulating Rad51 ubiquitination, AGS or GES-1 cells were co-transfected with plasmids expressing p62, Rad51 and ubiquitin, and then treated with a proteasomal inhibitor MG132. We found that overexpression of p62 resulted in an increase in the polyubiquitination of Rad51, compared with transfection controls (Figure 8E). Moreover, Cycloheximide chase assay showed that the knockdown of p62 by shRNA significantly extended the half-life of Rad51 protein (Figure 8F). This effect is likely attributable to the interaction of p62, ubiquitin and Rad51 as predicted in the STRING database (Figure S3B). Because the UBA domain of p62 is required for its binding to an ubiquitinated substrate, the plasmid expressing p62-ΔUBA was generated by deleting its C-terminal UBA domain. Rad51 was ubiquitinated on by intact p62, but not p62ΔUBA, indicating that p62 UBA domain is essential for p62-mediated Rad51 ubiquitination (Figure 8G). Taken together, these findings indicate that *H. pylori* infection induces the accumulation of p62, which promoted Rad51 ubiquitination and degradation via through the direct interaction between its UBA domain and Rad51.

### Expression profiles of p62 and Rad51 over the histopathologic cascade of human gastric cancer

To examine relevance of our in vitro and in vivo findings to clinical manifestations, CNAG, IM to Dys, GC from human subjects were collected and evaluated for the expression profiles of p62 and Rad51. Levels p62 were progressively elevated in the gastric tissues from CNAG, IM to Dys. However, p62 level was slightly lower in GC tissues than in Dys, although it was still significantly higher compared to CNAG group (Figure 9A). Interestingly, we found that the levels of Rad51 were significantly increased in Dys and GC (the severe precancerous and cancerous lesions) than in CNAG and IM (the mild and moderate lesions) (Figure 9B). These findings suggested that multiple factors, not just *H. pylori* infection, were involved in Rad51 regulation and genome integrity in gastric tumorigenesis. To further characterize the effect of *H. pylori* infection on their expression, gastric tissues were divided into *H. pylori*+ and *H. pylori*− groups. There were significantly decreased levels of Rad51 in *H. pylori*-infected Dys tissues, as compared to *H. pylori* non-infected Dys tissues. In addition, levels of p62 were significantly higher in *H. pylori*+ IM tissues compared to *H. pylori*− IM tissues (Figure 9C). These data indicate that *H. pylori* infection could promote elevation of p62 and reduction of Rad51 in the select histopathologic stages of human gastric cancer.

**Figure 9.**
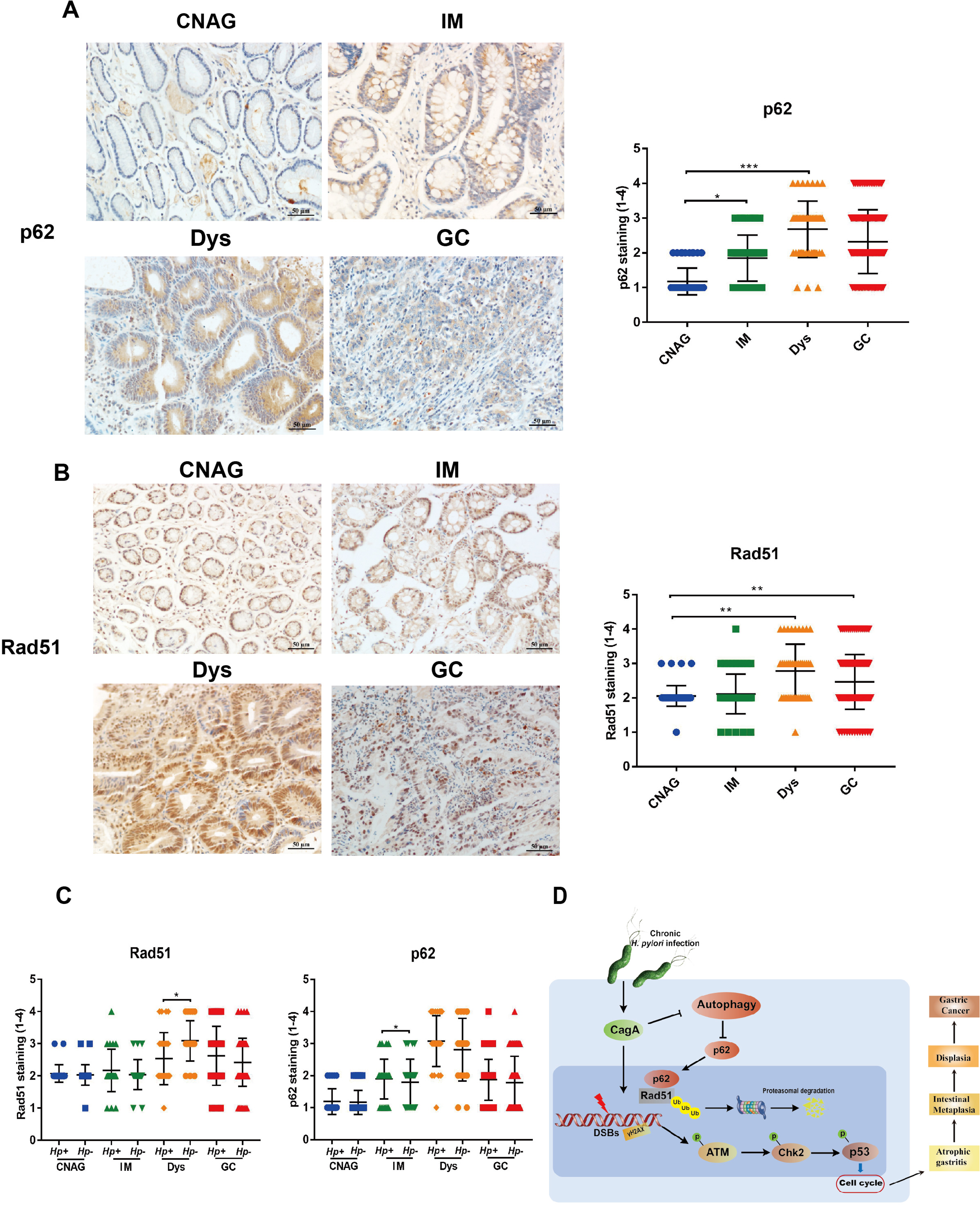
The expression of p62 and Rad51 in the human gastric carcinogenesis cascade. (A, B) Immunohistochemical staining for p62 and Rad51 in serial tissue sections from human gastric mucosa with different stages (CNAG, IM, Dys and GC) of the Correa histopathological cascade. (C) Expression levels of p62 and Rad51 were compared between *H. pylori*+ and *H. pylori*-gastric tissues with different stages (CNAG, IM, Dys and GC) of the Correa histopathological cascade. (D) Models of the regulation of autophagy-DNA damage axis in response to *H. pylori* infection. \*\**P*<0.01, \*\*\**P*<0.001.

## Discussion

*H. pylori* has been recognized to be one of the most common human pathogens. Epidemiological data suggest that approximately 90% of gastric cancer are attributable to *H. pylori* infection(Moss, 2017). Genomic instability is one of cancer hallmarks(Hanahan & Weinberg, 2011). Recently, DNA damage has been linked to autophagy, an intracellular degradation process by which cytoplasmic materials are delivered to the lysosome for digestion(Eliopoulos et al., 2016). Despite that autophagy has been implicated in *H. pylori* pathogenesis(Greenfield & Jones, 2013), the mechanism is still lacking. In this study, in vivo and in vitro data showed that *H. pylori* infection induced DSBs, triggered activation of DDR and impaired DNA repair via reduction of the DNA repair proteinRad51; this process is apparently CagA-dependent *in vitro.* Such reduction of Rad51 was mediated by H. pylori-induced suppression of autophagy, which led to accumulation of the autophagic substrate p62. Subsequently, the accumulation of p62 resulted in ubiquitination and degradation of Rad51 through the interaction of the UBA domain of p62 with Rad51. These findings are supported by elevation of p62 in concert with downregulation of Rad51 noted in human stomach samples with advanced precancerous gastric lesions. Based on these findings, a working model is proposed to depict the molecular mechanism underlying elevation of *H. pylori* CagA-induced DSBs (Figure 9D).

Cellular DNA damage can be caused by exposure to exogenous genotoxic agents. DSBs is a major cytotoxic lesion and must be repaired properly to maintain chromosomal integrity. Our data demonstrated that carcinogenic *H. pylori* can activate DNA damage response signalling, including induction of DSBs assessed by γH2AX, activation of checkpoint response by phosphorylation of ATM, Chk2 and p53. As a result, infection with *H. pylori* induced S cell cycle arrest, suggesting the rate of DNA synthesis has been decreased. HR and NHEJ are two main pathways to repair DSBs [36]. Our previous studies have shown that *H. pylori* infection is correlated with the levels of NHEJ marker Ku70/80[28]. In this study, we found that H. pylori stimulation induces the expression of Rad51 that is crucial for HR repair. However, upon exposure to prolonged infection of *H. pylori* (at 12h or 24h), DNA repair was impaired due to the loss of repair protein Rad51. As a result, which could potentially lead to genome instability. Such effect could lead to an increase of mutation rates and is associated with tumorigenic potential (Tubbs & Nussenzweig, 2017). The *H. pylori*-induced loss of Rad51 supports previous reports showing that the effects of genomic instability were induced by *H. pylori* infection (Koeppel, Garcia-Alcalde et al., 2015). It is worth nothing that Rad51 was overexpressed in human GC tissues compared with CNAG tissues, which is apparently inconsistent with H. pylori-modulated expression patterns of Rad51noted in gastric epithelial cells, gerbils or C57BL/6 mice. This discrepancy may result from the findings that Rad51 expression in human gastric tissues is regulated by multiple factors. It has been reported that elevated Rad51 was observed in a wide range of human tumors(Gachechiladze et al., 2017). On one hand, upregulation of Rad51 could compensate for the homologous recombination defects in another repair genes-deficient (*e. g.* BRCA1/2) tumors. On the other hand, overexpression of Rad51 in tumor cells is responsible for radio- and chemotherapeutic resistance through excessive DNA repair mechanism(Ward, Khanna et al., 2015).

It has been documented that that initial *H. pylori* infection transiently induced autophagy, which may be a host defence mechanism to mitigate toxin-induced damage(Hu, Zhang et al., 2019). And VacA was found to be sufficient to induce autophagosome formation(Terebiznik, Raju et al., 2009). Consistent with these reports, our in vitro studies showed that autophagy was induced following *H. pylori* infection at 6HPI, whereas prolonged exposure to *H. pylori* infection resulted in loss of autophagy in both gastric epithelial cells at 24 HPI and gerbils at 6 MPI. The virulence factor CagA played an important role in this process. This finding is supported by several previous studies reporting that persistent *H. pylori* CagA negatively regulated autophagy and promoted chronic inflammation (Li et al., 2017, Muhammad, Nanjo et al., 2017). It is plausible that persistent *H. pylori*-disrupted autophagy may facilitate accumulation of genotoxic products, which thereby contributing to gastric tumorigenesis.

A growing evidence has linked autophagy to DNA damage response. Autophagy can be activated by diverse DNA damage factors such as radiation, oxidative stress and chemicals. It has been reported that various DNA damage regulators including ATM, PARP1, and p53 are associated with rapid induction of autophagy, which in turn modulate DNA repair pathways. However, when DNA damage is beyond repair, autophagy is typical for the initiation of cell apoptosis and cell death(Hewitt & Korolchuk, 2017, Roos et al., 2016). In our study, our data indicate that inhibition of autophagy aggravated DNA damage via suppression of Rad51 in *H. pylori*-infected gastric epithelial cells. Treatment with autophagy inducer rapamycin or starvation induced G2/M cell cycle arrest, which prevented G2/M cells from entering mitosis. These findings suggest that dysregulated DNA damage response to *H. pylori* infection is attributed to altered cell cycle progression. This premise is supported by previous data the interaction between autophagy and regulation of cell cycle(Zheng, He et al., 2019). Taken together, we hypothesize that early infection with *H. pylori* induced DSBs, which can be sufficiently repaired by concurrently induced autophagy. However, long-term *H. pylori* infection can lead to loss of autophagy, which led to accumulation of unrepair DSBs through inhibition of the DNA repair protein Rad51.

It has been reported that the selective autophagy adaptor p62 could connect two biological processes: autophagy and DNA repair. P62 accumulation can result in inhibition of DNA-damage-induced histone H2A ubiquitination through the regulation of RNF168(Wang, Zhang et al., 2016). Also, it has been found that p62 suppressed Rad51-mediated repair mechanism by proteasomal degradation of FLNA(Hewitt et al., 2016). Our data indicate: (1) *H. pylori* infection upregulated p62 and downregulated Rad51 protein levels in vitro and in vivo models; (2) Transient overexpression of p62 apparently suppressed Rad51 expression, whereas knockdown of p62 increased Rad51 levels; (3) we found that p62 directly interacts with Rad51; (4) overexpression of p62 was associated with increased Rad51 ubiquitination, knockdown of p62 significantly prolonged the half-life of Rad51, and p62 induced Rad51 ubiquitination via the direct interaction of its UBA domain with Rad51. Based on our findings, we proposed that accumulation of p62 caused by *H. pylori* induced Rad51 ubiquitination and proteasomal degradation, which resulted in impairment of DSBs repair and genome instability.

In summary, our findings presented in this study provide new mechanistic insights into the relationship between autophagy and DNA damage response in *H. pylori*-associated gastric carcinogenesis. Chronic *H. pylori* infection induces the loss of autophagy and subsequent accumulation of autophagic substrate p62. The accumulated p62 results in ubiquitination of Rad51 via the direct interaction of its UBA with Rad51, thereby suppressing capability of the DNA damage repair. These cellular events allow *H. pylori* to elevate DSBs and genomic instability which potentially promotes gastric carcinogenesis. Further elucidation of this molecular mechanism will shed new insight into the development of more effective therapeutic strategies for preventing and curing *H. pylori* infection-associated gastric diseases.

## Materials and Methods

### Cell culture and *H. pylori* strains

Human gastric cancer AGS cells were cultured with DMEM/F12 (Gibco, CA, USA) containing 10% fetal bovine serum (FBS) and 1% penicillin/streptomycin (Gibco). The immortalized human gastric GES-1 epithelial cells were cultured in RPMI 1640 (Gibco) with 10% FBS and 1% penicillin/streptomycin. Cells were incubated at 37°C under a 5% CO2 atmosphere.

Previously characterized *H. pylori* strain CagA^+^ATCC43504 (NCTC11637) was used in this study(Crabtree, Farmery et al., 1994). Additionally, *H. pylori* strain 7.13 and its isogenic *cagA* mutant were kindly provided by Dr. Richard Peek at Vanderbilt University Medical Centre, Nashville, TN, USA. All *H. pylori* strains were cultured on Campylobacter agar plates containing 10% sheep serum and incubated at 37°C under microaerophilic conditions. The bacteria were suspended in DMEM/F12 or RPMI 1640 and the concentration were estimated by spectrophotometry (OD_600nm_). Subsequently cells were supplemented with fresh medium without antibiotic, followed by incubation with *H. pylori* at specified time and different multiplicities of infection (MOIs).

### Animals and infections

All experiments and procedures carried out on animals were approved by the Ethics Committee of The First Affiliated Hospital of Nanchang University. Specific pathogen free (SPF) male Mongolian gerbils were challenged with *H. pylori* by previously described methods (Yang, Xie et al., 2015). In addition, five to six-week-old male SPF C57BL/6 mice (Hunan SJA Laboratory Animal Co Ltd, Hunan, China) were administrated with 2×10^8^ colony forming units (CFU)/ml bacterial suspension of *H. pylori* 43504 once every other day for 5 times. After infection with *H. pylori* for 3 months, mice were given with autophagy inducer everolimus (1.5mg/kg, every other day) for a total of 6 times.

### Histopathology

Then mice were euthanized and linear strips of gastric tissues extending from the squamocolumnar junction through proximal duodenum were collected. Colonization of *H. pylori* of animals was examined by immunohistochemistry or fluorescence in situ hybridization (FISH) staining. H&E staining was used to evaluate gastric pathology. Gastric lesions were graded on a scale from 0 to 4 as the extent of inflammatory cells infiltration in the mucosa and submucosa(Ge, Sheh et al., 2018).

### Gastric specimens and Immunohistochemistry

Human gastric tissue samples were obtained from patients undergoing gastroduodenoscopy or gastrectomy at the First Affiliated Hospital of Nanchang University. Embedded-paraffin tissues were collected as described previously(Xie et al., 2014), which included 56 of CG, 53 of IM, 47 of Dys, 146 of GC. The study was approved by the Ethics Committee of The First Affiliated Hospital of Nanchang University. Immunohistochemistry (IHC) was performed as previously described(Li, Feng et al., 2018). Briefly, the slides were deparaffinized, rehydrated. Endogenous peroxidase activity was blocked in 3% H_2_O_2_, and antigens retrieval utilizing microwave oven heating. Tissue sections were incubated with primary antibodies overnight at 4°C, and followed by incubation with secondary antibody. Finally, slides were developed in diaminobenzidine (DAB) solution, and counterstained with haematoxylin and mounted with coverslips.

### Antibodies, siRNA, plasmids and reagents

Antibodies and their sources were as follows: Anti-γH2AX (05-636), Anti-Rad51 from Millipore, Anti-ATM (ab78), p-ATM (S1981) (ab36810), Chk2 (ab47433), p-Chk2 (ab85743), BRCA2 (ab27976), p-P53 (S15) (ab1431), p-P53 (S20) (ab59206) from Abcam (Cambridge, MA, USA); LC3B (3868S), and GAPDH (2118S) from Cell signalling Technology (Beverly, MA, USA). Goat anti-Rabbit and anti-mouse secondary antibodies were purchased from Invitrogen (Thermo Fisher Scientific, Suwanee, GA, USA). For immunofluorescence assay, 4’,6-diamidino-2-phenylindole (DAPI), anti-rabbit or anti-mouse conjugated to Alexa Fluor 488, anti-rabbit or anti-mouse conjugated to Alexa Fluor 594 were from Invitrogen.

Pharmacological agents: rapamycin (R8781), everolimus (SML2282), 3-methyladenine (3-MA) (M9281), Bafilomycin A1 (BafA1) (B1793), Cycloheximide (C7698) were obtained from Sigma (St. Louis, MO, USA). Proteasomal inhibitor MG132 was purchased from Selleck Chemicals.

The recombinant plasmid of pcDNA3.0 HA-Rad51 were kindly provided by Dr. Xingzhi Xu at Capital Normal University. Myc-p62, Flag-Rad51, HA-Ubiquitin plasmids and Rad51 siRNA, p62 shRNA were purchased from Gene-Chem, Shanghai, China.

### Western blotting

Tissue or cells were lysed in lysis buffer with protease inhibitor cocktail (Roche, Amherst, CA, USA). Protein concentrations were determined by the BCA assay kit (Thermo Fisher Scientific). Protein samples (25μg) were separated by SDS-PAGE and transferred to nitrocellulose membranes. PageRuler Prestained Protein Ladder (cat. #26616) was obtained from Thermo Fisher Scientific (Waltham, MA, USA). The membranes were blocked in 5% blocking buffer (5% non-fat dry milk in TBST buffer) at room temperature for 1h and then incubated with primary antibodies overnight at 4°C. The membranes were then incubated with secondary antibodies at room temperature for 1h. The protein bands were imaged and quantified by using the the BioRad-ChemiDoc XR+ system. The signal intensity of each target protein band was normalized to GAPDH.

### Immunofluorescence

For immunofluorescence, human gastric epithelial cells were fixed with 4% formaldehyde in PBS for 15 min at room temperature. The cells were permeabilized with 0.25% Triton X-100 for 15min and then blocked in PBS with 3% BSA for 1h. Subsequently, the cells were incubated with primary antibody and then with a secondary fluorescent antibody. Cell nuclei were counter-stained with DAPI. All slides were evaluated using a fluorescent microscope (Nikon C2).

### Flow cytometry

Cell pellets were resuspended in Buffer A (1mg/ml Sodium citrate, 0.1%Triton-X-100, 20μg/ml RNase A and 100μg/ml propidium iodide) at 4°C for 30min in the dark, and then were analysed by a flow cytometer (BD Accuri C6).

### Comet assay

The comet assay was performed as previously described(Xie et al., 2014). Briefly, cells were suspended in low-melting-point agarose and spread on glass slide. The slides were incubated in lysis buffer at 4°C for 1h and then were electrophoresed at 30V for 30min. Comet tails were stained with propidium iodide (PI). More than 70 cells were analysed under the Nikon C2 confocal microscope.

### Transmission electron microscopy

The gastric cells were cocultured with *H. pylori* for indicated time. Cells were fixed in glutaraldehyde for 1.5h at 4°C, then washed and fixed again in 1% OsO4, and dehydrated in graded ethanol and embedded in Epon-Araldite resin. Ultrathin sections were cut and stained with uranyl acetate and lead citrate, and visualized under a HT7700-SS electron microscope (HITACHI, Japan).

### Real-time quantitative PCR analysis

Transcript levels of IL-1β, IL-6, IL-8, IL-10, IL-17 and Rad51 were assayed using qPCR as previously methods(Li et al., 2018). The following primers were used: mouse IL-1β, forward primer 5’-GACCTTCCAGGATGAGGACA-3’, reverse primer 5’-AGGCCACAGGTATTTTGTCG; mouse IL-6, forward primer 5’-TCTCCAGCAACGAGGAGAAT-3’, reverse primer 5’-TGTGATCTGAAACCTGCTGC-3’; mouse IL-10, forward primer 5’-AAGCTGAGAACCAAGACCCAGAC-3’, reverse primer 5’-AGCTATCCCAGAGCCCCAGATCCGA-3’; mouse IL-17, forward primer 5’-ACCTCAACCGTTCCACGTCA-3’, reverse primer 5’-CAGGGTCTTCATTGCGGTG-3’; mouse GAPDH, forward primer 5’-GTAGCAAAGGGAATGGGTCT-3’, reverse primer 5’-AGATGGTGAAGGGCTAATGG-3’; human Rad51, forward primer 5’-CTATGTAGCAAAGGGAATGGG-3’, reverse primer 5’-AAGCAGGTAGATGGTGAAGG-3’; human GAPDH, forward primer 5’-GTAGCAAAGGGAATGGGTCT-3’, reverse primer 5’-AGATGGTGAAGGGCTAATGG-3’.

### Cycloheximide chase assay

Cycloheximide chase assay was performed to characterize Rad51 protein stability. Briefly, AGS cells were transfected with p62 shRNA plasmid and then treated with cycloheximide (50ng/ml). Rad51 protein levels were examined at specified times.

### Immunoprecipitation assay

After transfected with expressing p62 or Rad51 plasmid, cells were collected and lysed with RIPA lysis buffer. All samples were processed using the Immunoprecipitation Kit (Abcam, ab206996) according to the manufacture recommendations. In brief, lysed cells were incubated with a primary antibody overnight at 4°C. Prior to immunoprecipitation, protein A/G beads were washed with the wash buffer. After antibody binding, p62 or Rad51 was immunoprecipitated with agarose beads for 2h at 4°C. Western blotting was performed to detect the levels of immunoprecipitated proteins.

### In vitro ubiquitination assay

AGS cells were transfected with plasmids (Flag-Rad51 or Myc-p62) and ubiquitin for 48h, then co-cultured with *H. pylori* strain and treated with MG132 for 6h. Immunoprecipitation assay was performed as described(Hu, Liu et al., 2019).

### FISH assay

FISH assay was performed to detect *H. pylori* colonization in animal paraffin-embedded animal gastric biopsy specimens using a *H. pylori*-specific fluorescent probe-Hpy (5’-CACACCTGACTGACTATCCCG-3’). The probe labelled with fluorochrome Cy3 at 5’-end were synthesized by Genscript (shanghai, China). The specimens were overlaid with a hybridization buffer (0.9 M NaCl, 20mM Tris-HCl [pH 7.2], 0.01% sodium dodecyl sulfate) containing 30% formamide and an Hpy oligonucleotide probe (10ng/μl). Hybridization was performed at 48°C overnight in a humid chamber. Then, the slides were washed in the pre-warmed washing buffer I (0.9 M NaCl, 20 mM Tris-HCl [pH 7.2], and 0.01% sodium dodecyl sulfate) and the washing buffer II (0.9 M NaCl, 20 mM Tris-HCl [pH 7.2]) for 15mins, respectively. Subsequently, the slides were stained with DAPI and examined under the fluorescent microscope (Nikon C2).

### Statistical assay

All data were presented as mean ± standard error of mean (S.E.M). All statistical analyses were performed using SPSS 20.0 software. Mann-Whitney, Student’s t-test and one-way analysis of variance (ANOVA) were applied depending on the data set. *P*<0.05 (***, *P* < 0.001, **, *P* < 0.01, *, *P* < 0.05) were considered significant.

## Acknowledgments

We thank Prof. Zhongming Ge (Massachusetts Institute of Technology, Cambridge, MA, USA) for editing of the manuscript. We also thank Prof. Richard Peek (Vanderbilt University Medical Centre, Nashville, TN, USA) for kindly providing the H. pylori strain 7.13 and its CagA-mutant. We also thank Dr. Xingzhi Xu (Capital Normal University, Beijing, China) for pcDNA3.0 HA-Rad51 plasmid.

## Author Contributions

CX, HW, NSL, NHL conceived, designed the study and analysed data. NSL and DJW drafting the manuscript. YH, CP and JC collected human specimens and analysed immunohistochemical data.CH, DJW, SX, YZ and YX provided technical support. CX and NHL Obtained funding support. NHL supervised and oversaw the study. All the authors critically revised the manuscript and provided intellectual content.

## Ethics approval

The biopsy specimens were obtained under protocols and all animal experiments were approved by the ethics committees of The First Affiliated Hospital of Nanchang University.

---

**Figure S1.** (A) Representative immunohistochemical staining for ATM and Chk2 in a series of human gastric tissues with CNAG, IM, Dys and GC. (B) Quantitative analysis of the immunohistochemical staining results in gastric tissues.

**Figure S2.** (A) Immunohistochemistry staining analysis and (B) FISH staining analysis showing *H. pylori* colonization of gastric mucosa of mouse models. (C) Q-PCR analysis showing the levels of inflammatory response factors including IL-1β, IL-10, IL-6 and IL-17 in gastric mucosa of mouse models.

**Figure S3.** (A) Q-PCR analysis showing Rad51 mRNA levels in gastric cells following *H. pylori* infection at different time points. (B) The interaction between Rad51 and p62 protein was predicted through STRING database.

